# KoNA: Korean Nucleotide Archive as a New Data Repository for Nucleotide Sequence Data

**DOI:** 10.1101/2023.07.27.550450

**Authors:** Gunhwan Ko, Jae Ho Lee, Young Mi Sim, Wangho Song, Byung-Ha Yoon, Iksu Byeon, Bang Hyuck Lee, Sang-Ok Kim, Jinhyuk Choi, Insoo Jang, Hyerin Kim, Jin Ok Yang, Kiwon Jang, Sora Kim, Jong-Hwan Kim, Jongbum Jeon, Jaeeun Jung, Seungwoo Hwang, Ji-Hwan Park, Pan-Gyu Kim, Seon-Young Kim, Byungwook Lee

## Abstract

During the last decade, generation and accumulation of petabase-scale high-throughput sequencing data have resulted in ethical and technical challenges, including access to human data, and transfer, storage, and sharing of enormous amount of data. To promote data-driven research in biology, the Korean government announced that all the biological data generated from government-funded research projects should be deposited in the Korea BioData Station (K-BDS), which consists of multiple databases for individual data types. We introduce the Korean Nucleotide Archive (KoNA), a repository for nucleotide sequence data. As of July 2022, the Korean Read Archive in KoNA has collected over 477 TB of raw next generation sequencing data from several national genome projects. To ensure data quality and prepare for international alignment, a standard operating procedure (SOP) was adopted, which is similar to the International Nucleotide Sequence Database Collaboration. The SOP includes quality control processes for submitted data and metadata using an automated pipeline followed by manual examination. To ensure fast and stable data transfer, a high-speed transmission system called GBox is used in KoNA. Furthermore, the data uploaded to or downloaded from KoNA through GBox can be readily processed in a cloud-computing service for genomic data analysis called Bio-Express. This seamless coupling of KoNA, GBox, and Bio-Express enhances data experience including submission, access, and analysis of raw nucleotide sequences. KoNA not only satisfies the unmet needs for a national sequence repository in Korea, but also provides datasets to researchers globally and contribute to advances in genomics. KoNA is available at https://www.kobic.re.kr/kona/.

## Introduction

The drastic price decrease in next-generation sequencing (NGS) data and improvement in sequencing platforms with higher data throughput produce more than tens of petabases of raw sequencing data [1,2]. For example, in Korea, a few petabases of NGS data in diverse biological systems have been generated from several national projects, such as the Korea Post-Genome Project. To deposit, store, and share the enormous amount of raw nucleotide sequence data, there have been many efforts to construct repository databases by national and international data centers, such as the National Center for Biotechnology Information (NCBI), the European Molecular Biology Laboratory’s European Bioinformatics Institute (EMBL-EBI), the DNA Data Bank of Japan (DDBJ), and the National Genomics Data Center (NGDC) [3−6]. These publicly available repositories provide datasets to global scientific communities and facilitate data reuse, which can reduce costs and usher advances in genomics. Despite the impressive contribution of these repositories, additional national repositories are emerging in many countries mainly for three reasons. First, many nationwide genome projects are under way in many countries, which are producing huge amount of data that need to be archived. Second, domestic network has a larger upload and download bandwidth and enables faster data transfer than international network. Third, deposition and access of human data are controlled by domestic laws due to the sensitive nature of human data [4,6]. The Korea BioData Station (K-BDS), which was developed by the Korea Bioinformation Center (KOBIC), is a data archive that was initiated to address these three reasons.

First, K-BDS serves as a central archive for all the biological data generated from Korean government-funded research projects. In 2020, the Korean government announced an initiative called the “Strategy on Biological Research Resource Big Data” to promote data-driven research in biology. The announcement of the initiative was followed by an introduction of a data policy that mandates all the biological data generated from government-funded research projects to be deposited in a central data archive, for which K-BDS was designated [7]. Given the data policy that is applied to all biological data types, K-BDS consists of several databases, each of which specialize in collecting and sharing a specific data type, such as NGS, microarray, quantitative proteomics, metabolomics, and bioimages. Second, K-BDS enables significantly faster submission and sharing of data within the nation. For domestic researchers in Korea, data submission and access to K-BDS (949 Mbps upload and 951 Mbps download) are about nine times faster on average than those to NCBI (123 Mbps upload and 107 Mbps download), according to the Communication Service Quality Evaluation Results by the Korea Ministry of Science and ICT in 2019 (http://www.smartchoice.or.kr/smc/info/evaluateDownload.do). Third, K-BDS handles sensitive human data in a way that complies with relevant Korean laws and government policies, such as the Bioethics and BioSafety Act and the Guideline for the Use of Health and Medicine Data.

In this article, we introduce the Korean Nucleotide Archive (KoNA), the nucleotide sequence data-specific database in K-BDS, to fulfill the unmet needs for the efficient collection, management, and sharing of nucleotide sequence data in Korea. KoNA currently accepts NGS data submission, with a plan to extend toward other types of nucleotide sequences, such as assembled sequences and primer sequences. Currently, the Korean Read Archive (KRA) in KoNA contains NGS data that encompass various sequencing experiments, namely genomics, transcriptomics, epigenomics, and metagenomics, and diverse organisms as well, such as humans, animals, plants, and microorganisms. Furthermore, KoNA is coupled to a high-speed data transmission system called GBox, which is in turn coupled to a cloud-computing service for large-scale biological data analysis called Bio-Express [8,9]. As all these services are offered in English language, they can be used for all the researchers in the globe.

## Results

### Overview of KoNA

Designated by the “Act on the Acquisition, Management, and Utilization of Biological Research Resources,” KOBIC is a national center for collecting biological data generated by Korean government-funded research projects. In 2021, KOBIC established K-BDS, which consists of several repository databases that are specific to individual data types. One such database is KoNA, which collects and provides primary nucleotide sequence data, such as FASTQ or FASTA formatted files generated by diverse NGS techniques, along with rich metadata. To prepare for international alignment with existing public databases, the data standard and structure of the following repository databases were examined: NCBI Sequence Read Archive (SRA), EMBL-EBI European Nucleotide Archive (ENA), DDBJ, and NGDC GSA [3−6].

Upon the examination of the existing repositories, we adopted the overall data structure implemented in those databases, which is used to hierarchically organize various metadata underlying the NGS data. In specific, we adopted the BioProject and the BioSample to capture descriptive information of research projects and biological samples, the Experiment to capture instrument and library information, and the Run to capture sequence data file. For the BioSample, we used only the INSDC’s Standard Packages but not the MIxS Packages of Genome Standards Consortium (GSC) yet.

Currently, KoNA only contains KRA, a sequencing read data archive that is akin to NCBI SRA and EMBL-EBI/ENA SRA, DDBJ Sequence Read Archive (DRA), and NGDC/GSA China Read Archive (CRA). KoNA is also systematically linked with the Korean Genome-Phenome Archive (KGA), a repository of controlled-access data for human genotypes and phenotypes, which is similar to the European Genome-Phenome Archive (EGA), the database of Genotypes and Phenotypes (dbGaP), and the Japanese Genotype-phenotype Archive (JGA) (**Figure 1A**). While most data deposited in KRA can be accessed without any restrictions, human-related data in KRA are only accessible after approval of data access request to KGA operated by the Korea Human BioData Bank.

**Figure 1.**
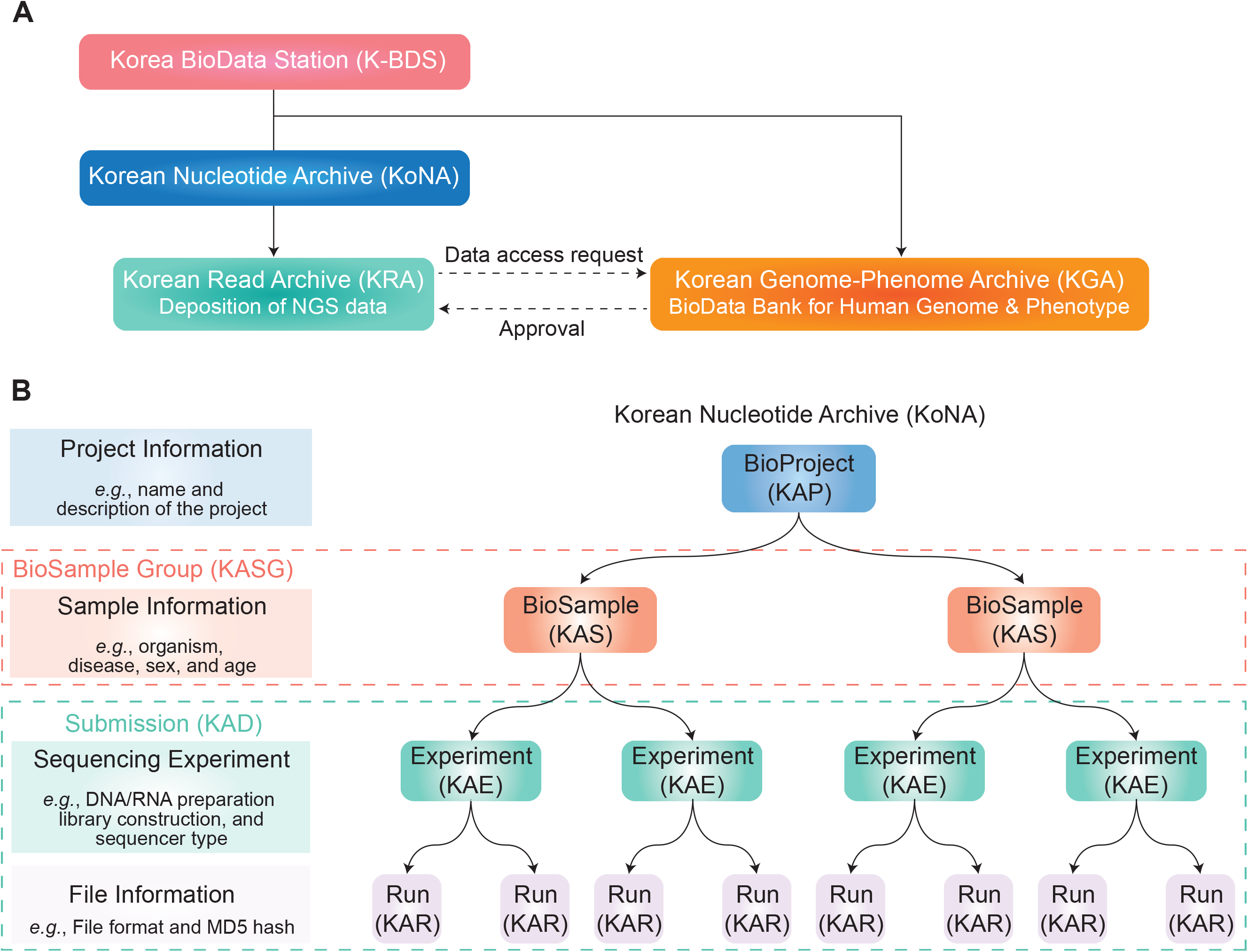
Database structure and data model of K-BDS and KoNA. **A**. Korea BioData Station (K-BDS) and Korean Nucleotide Archive (KoNA) database structure. KoNA, the national nucleotide sequence repository, is one of the databases in K-BDS. KoNA contains the Korean Read Archive (KRA), a primary NGS data repository. Furthermore, for human data, KoNA is systematically linked with the Korean Genome-Phenome Archive (KGA) of K-BDS, a database for human nucleotide sequences. All the primary NGS data of human samples are accessible in KRA after KGA approves data access request by users. **B**. The current data model of KoNA. It consists of four objects: BioProject, BioSample, Experiment, and Run. For each data object, description and attribute example, and prefix of accession ID are presented in the left panel and parentheses. The dashed boxes indicate groups of BioSamples (“BioSample Group”) and Experiments and Runs (“Submission”) with “KASG” and “KAD” as the prefixes for their accession IDs, respectively. Notably, the “K” in the prefixes of all the accession IDs denotes “Korea/Korean.

### Data model and structure of KoNA

The metadata model of KRA is equivalent to those established by the INSDC databases and the NGDC GSA, in which all the attributes are categorized into four objects (**Figure 1B**): study (i.e., “BioProject” object with KAP prefix; e.g., KAP210001), sample information (i.e., “BioSample” object with KAS prefix; e.g., KAS21000001), sequencing experiment (i.e., “Experiment” object with KAE prefix; e.g., KAE21000001), and data files (i.e., “Run” object with KAR prefix; e.g., KAR21000001). For the numerical part, the first two decimal digits indicate the year of accession ID issue, and the other decimal digits are sequential numbers assigned in chronological order. We denote a group of BioSamples as BioSample Group with KASG prefix (e.g., KASG210001) to manage and present relevant BioSamples in the system for user convenience. We issue an accession ID for a group of Experiments and Runs annotated in one submission batch (i.e., “Submission” object with KAD prefix; e.g., KAD2100001).

### Data submission and quality control

The user-friendly interface of the KoNA website helps users understand the data submission, retrieval, and access procedures intuitively and get the statistics of the data deposited in KoNA (e.g., the numbers of deposited BioProjects, BioSample Groups, Submissions, and Runs). To deposit data in KoNA, raw sequence data and accompanying metadata should be submitted through GBox and the website, respectively (**Figure 2**). Prior to the submission of metadata, primary raw sequence data, along with processed secondary data if available, should be uploaded to the user space in GBox. For primary data, KoNA allows FASTQ and FASTA file formats. Submission of FASTQ files is highly recommended when data were generated by short-read sequencing methods, such as Illumina and MGI platforms, to maintain the Phred quality scores for individual bases. For secondary data, KoNA allows file formats such as binary alignment map (BAM), sequence alignment map (SAM), variant call format (VCF), and tab-delimited text. Secondary data alone without any corresponding primary data are not accepted.

**Figure 2.**
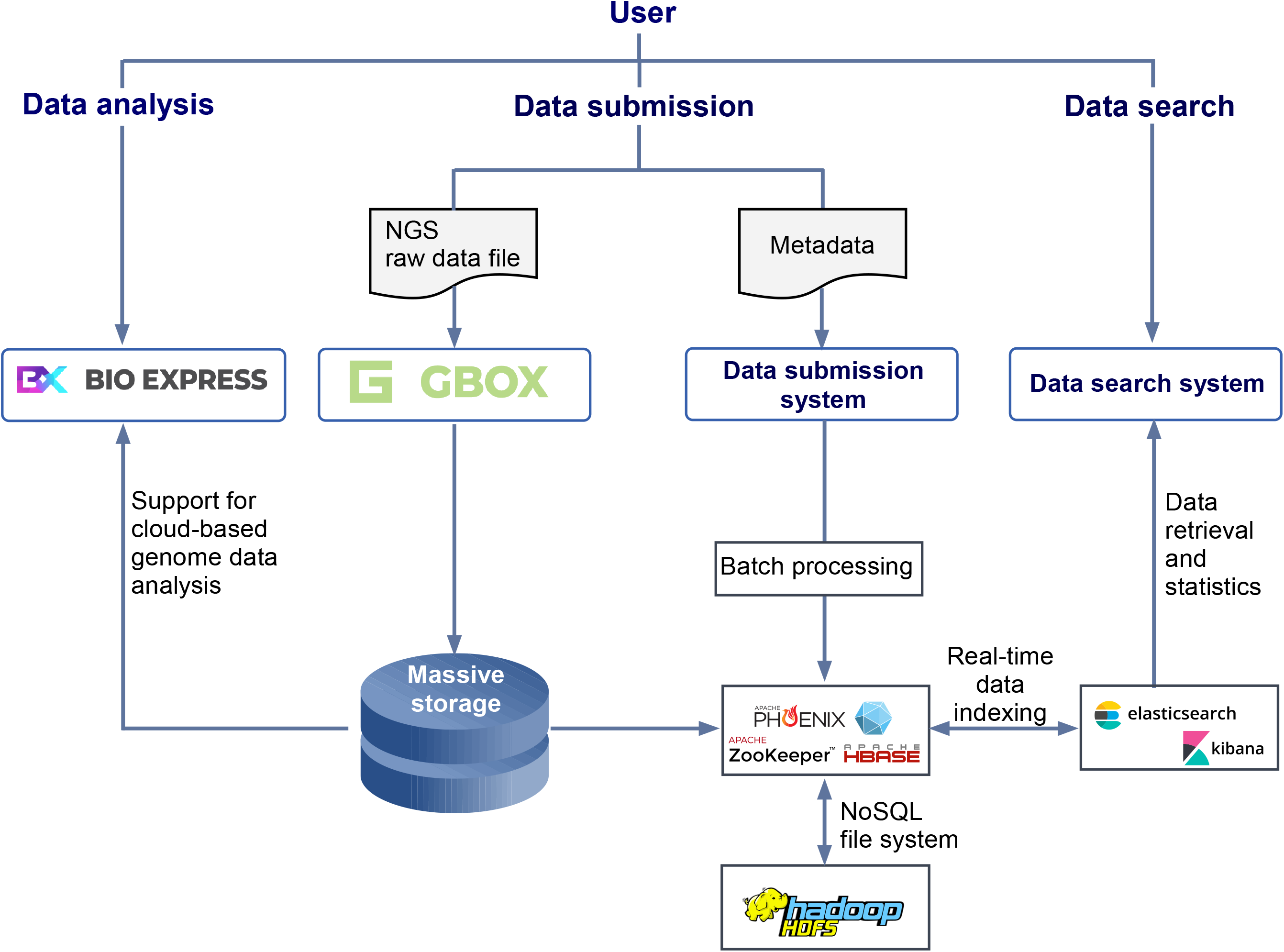
Workflow of data submission, quality control, and curation processes. During the submission process, the primary data (i.e., raw sequence data) and/or secondary data should be submitted through GBox, whereas the relevant metadata should be submitted on the website. During the quality control process, we confirm the MD5 hashes and file extensions of the submitted data and their metadata undergo automated and manual examination. Finally, all these data and metadata are migrated into KoNA, followed by the data curation process by KoNA curators.

As described in the submission guideline on the website, submission of metadata proceeds in the order of BioProject, BioSample Group, and Submission. Online spreadsheets with attribute names previously established by the INSDC are used to provide the details of multiple BioSamples in a BioSample Group, and those of multiple Experiments and Runs in a Submission. Moreover, to match the individual Experiments with the primary and secondary data uploaded in GBox, file names and directories in the GBox user space should be described in the Submission spreadsheet.

We aim to collect high quality data that can increase data usability. Therefore, we successively perform quality control for both primary and secondary data first, followed by metadata. For primary and secondary data, we first automatically validate MD5 hashes and confirm the file formats or extensions with those inputted to the corresponding metadata spreadsheet. For each primary data, the number of spots, reads, and bases are assessed by in-house scripts. On the other hand, the metadata submitted to the KoNA website are first subjected to an automated examination to verify the integrity of each attribute in the metadata. Currently, the automated examination includes checkup of whether mandatory fields were inputted, and whether correct variable type, such as numeric, character, and date, was inputted for each attribute. Further development is also underway to expand the functionality of the automated examination. The automated examination is followed by a manual examination and correction by KoNA curators, who are data experts in KOBIC. The curators communicate closely with submitters, examine the research context, and correct the metadata to fill in all the critical contents required for data reanalysis and for conveying the original biological contexts of the submitted data to data users. Upon completion of the quality control processes, the submitted data are migrated into KoNA and the KRA database with accession IDs for individual objects. Finally, KoNA curators group the deposited studies into data collections based on their research contexts during data curation processes. All these curated collections are to be shown in secondary databases in KoNA. To aid data submission, user documentations in English is provided, and helpdesk is available as well, whose contact information is listed in the Contact Us webpage.

### Data retrieval and access

For data deposited in KoNA, all publicly accessible BioProjects are listed in the “Browse” menu. Moreover, users can search relevant BioProjects, BioSample Groups, and Submissions by keyword queries. To enable fast and flexible responses to a given query, we applied a search engine for big data based on elastic search [10] and developed a search engine optimized for the KoNA system. The KoNA system exhibits the following advantages. First, it stores the indexed data and is fully searchable in near real time. Second, it supports high-speed search for structured and unstructured data using a big data platform capable of parallel distributed processing. Third, it supports flexible extension of diverse data types using schemaless NoSQL. Fourth, it supports diverse programmatic access methods by supplying RESTful API. Finally, it visualizes real-time statistics from access logs using elastic search and Kibana tools.

Among a total of 666 Submissions deposited in the KRA database, 602 Submissions (90.4%) have been released as of July 2022. Journal reviewers and editors can access embargoed Submissions if the submitters issue access tokens for their data in the KoNA website. Users authorized by KGA can access 117 released Submissions out of a total of 137 controlled-access Submissions. The data request and access procedures in the KGA are in accordance with Korea’s “Bioethics and BioSafety Act” and related ethical restrictions. Submissions with nonhuman data are readily accessible to public.

### Data transfer and cloud-based data processing utilities

KoNA was designed as a package deal, in which users not only deposit and provide their nucleotide sequence data but also enhance data experience by tying KoNA together with a fast and reliable data transfer system and diverse genome analysis utilities. To this end, KoNA adopts a high-speed data transmission system called GBox, which can transfer massive sequence data ten times faster than the speed of the normal file transfer protocol (FTP) and HyperText Transfer Protocol (HTTP) [9]. Furthermore, uploaded or downloaded sequence data by GBox can be readily processed using data processing tools implemented with graphical user interface in a cloud-computing service for large-scale biological data analysis called Bio-Express. This cloud-computing service is equipped with data analysis utilities for various sequence data types, such as genome, transcriptome, epigenome, metagenome, and single-cell genome [8,9]. This seamless coupling of KoNA, GBox, and Bio-Express, each of which has a user-friendly interface, facilitates data experience including submission, access, and reuse of raw nucleotide sequences. Current quota per user of Bio-Express is 100 CPU core, 400 GB memory, and 1 TB storage.

### Data statistics and contents

As of July 2022, a total of 360 BioProjects, 653 BioSample Groups, 27,209 BioSamples, 666 Submissions, 27,634 Experiments, and 30,423 Runs are deposited in KoNA and KRA. The total 666 Submissions in the KRA database contain 477 TB of raw sequence data. Currently, KRA has less volume of deposited data than SRA (about 20 PB on September 2021) [1], or GSA and GSA-Human (about 9 PB in June 2021) [11]. However, the data volume in KRA is expected to be comparable to that of the other databases in upcoming years, as KoNA collects a substantial portion of raw sequence data generated from Korean government-funded research projects.

Regarding the 360 BioProjects, more than 198 principal investigators have participated in data deposition. Furthermore, the 27,209 BioSamples encompass 436 organisms, which correspond to 280 microbial, 62 animal, and 93 plant species as well as one human species (*Homo sapiens*). Notably, 26,127 out of the 27,207 BioSamples (96.0%) belonged to the top 20 species with the largest number of BioSamples, and 19,788 (72.7%) and 4,932 (18.1%) BioSamples originated from humans and metagenomes, respectively (**Figure 3A**). A total of 19,788 human BioSamples were annotated with more than 87 diseases. Interestingly, among the top 20 diseases with the largest number of human BioSamples in KoNA, 17 out of 20 diseases were cancer diseases (**Figure 3B**). Such data, which are provided with rich metadata after quality control process by KoNA curators, may reveal the current research trend of Korean government-funded research projects.

**Figure 3.**
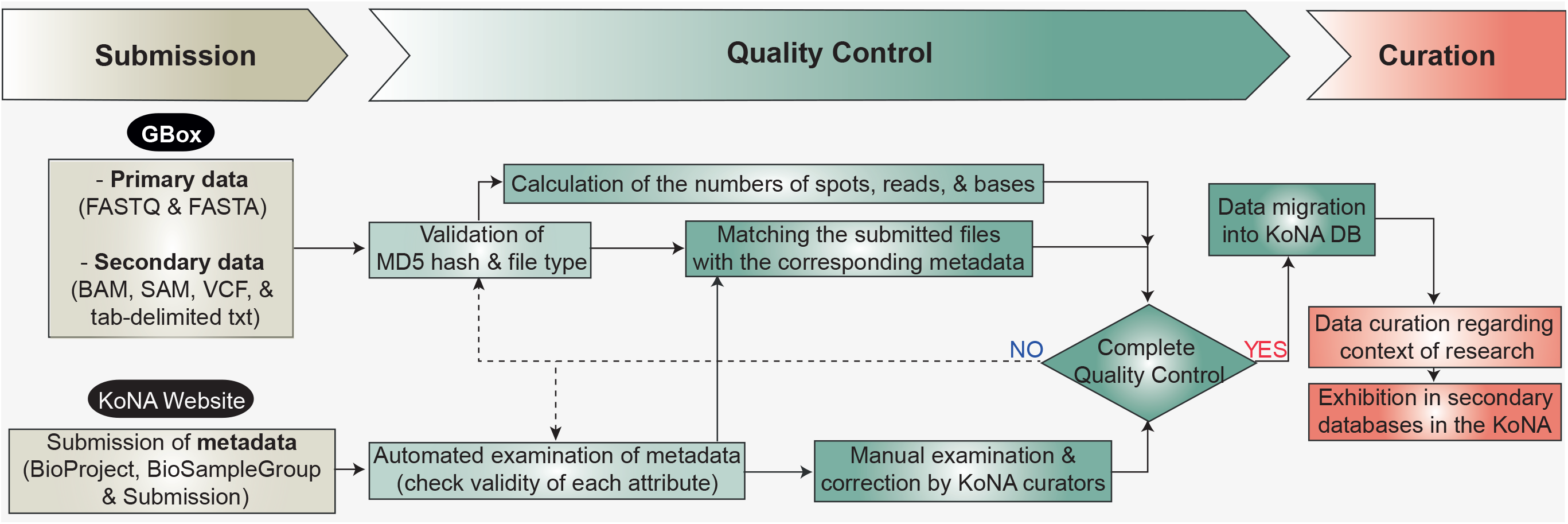
Summary of data statistics. **A**. Top 20 species in KoNA with respect to the number of BioSamples. Each color denotes the sample type (human, animal, microorganism, and plant) the species belongs to. **B**. The top 20 diseases in KoNA with respect to the number of human BioSamples. **C**. The number of strategies for generating raw NGS data. **D**. The number of sources used for sequencing experiments.

## Discussion

In this article, we introduced KoNA as the nucleotide sequence repository in K-BDS. Following the announcement of an initiative called the “Strategy on Biological Research Resource Big Data” by the Korean government in 2020, K-BDS was constructed to collect all types of biological data generated from government-funded research projects. KoNA is a subdivision of K-BDS specialized for nucleotide sequences. Currently KoNA accepts NGS data submission, with a plan to extend toward other types of nucleotide sequences, such as assembled sequences and primer sequences. We also plan to improve KGA in K-BDS, which currently has only a few functions for access request and approval of human data deposited in KRA. Because the data collection of KoNA is to be expanded to all primary nucleotide data, KGA is also updated to manage all those sequence data as well as the paired clinical data.

Although we have been successful in collecting and sharing domestic NGS data in Korea through KoNA over the years, we are also aware of the limitation of not yet being aligned with INSDC, the principal authority for international sharing of nucleic acid data. To overcome this limitation and progress into an internationally recognized data resource, we recently began an attempt to build close collaboration with INSDC by further strengthening our existing collaboration with DDBJ. In specific, an arrangement and preparation are underway for confirming the agreement of data models between KoNA and INSDC, and sharing the data in KoNA with INSDC through DDBJ.

To ensure quality of deposited data, data curation is performed by expert curators at KOBIC. They categorize the data and metadata based on the research context to increase data findability and accessibility. Related BioProjects, BioSample Groups, and Submissions are categorized into a collection based on research context. For example, whole-genome sequencing (WGS) data from human lung cancer patients are automatically collected in terms of “WGS”, “lung cancer”, and “*homo sapiens*”. The KoNA website is under development to show these collections with attribute information. Such data collection sets enable users to access the preferred sets deposited in KoNA and run advanced queries on the database by selecting from the previously extracted attribute information of the collection. Next, to enhance usability with the collection sets, we plan to construct secondary databases, such as 3DIV [12], iCSDB [13], ChimerDB [14], and GEMiCCL [15], which will provide useful information by using processed NGS and other nucleotide sequence. All these additional developments that are currently under way or planned will not only satisfy the unmet needs for a national sequence repository in Korea but also provide datasets to researchers globally and contribute to advances in genomics.

## Materials and Methods

### Establishment of the integrated service for genomic big data in KOBIC

To enhance data submission, transfer, retrieval, and reuse, an advanced integrated service for genomic big data was established by KOBIC through integrating Bio-Express, GBox, and NoSQL-based database architecture (**Figure 4**).

**Figure 4.**
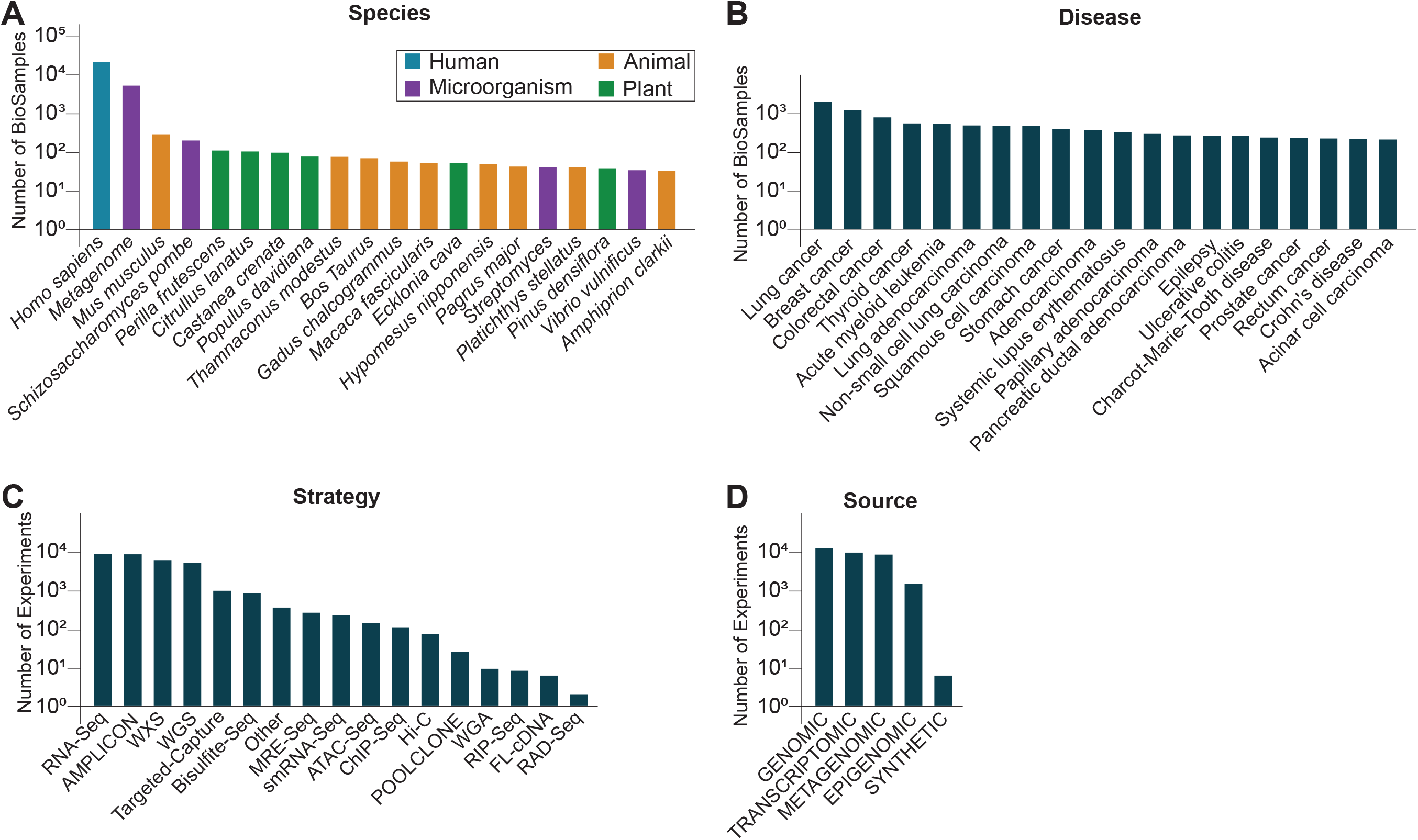
Schematic of the integrated service for genomic big data. The interconnection between KoNA, GBox, and Bio-Express, supported by the HDFS and NoSQL-based database and elasticsearch, improves user convenience in data submission, transfer, retrieval, and reuse.

GBox, which was developed by KOBIC, provides fast data transfer and high-capacity cloud storage on this integrated service. Two relay servers secure stability and enable about five times faster upload speed than Galaxy FTP (9.7 Mb/sec and 1.8 Mb/sec for GBox and Galaxy FTP, respectively). After data transmission with GBox, the data are divided into blocks and stored in the Apache Hadoop file system (HDFS; version 2.7.1) to facilitate high-capacity cloud storage.

Data reuse is facilitated by supporting analysis of genomic big data with Bio-Express, which is equipped with various bioinformatics analysis pipelines and uses the Hadoop-based distributed and parallel processing for fast analysis of big data. For example, for variant calling process of 500 GB WGS data with the GATK pipeline in the same hardware (Intel Xeon Gold 6132 CPU @ 2.60 Hz x 28 with 37 GB memory), the use of parallel computing in Bio-Express is about three times faster than an ordinary setting.

As the Backend database of the integrated service, the Apache HBase (version 2.0.5), a NoSQL database, with Apache Zookeeper (version 3.6.0) and Phoenix were adopted on the HDFS. This NoSQL-based database enables effective storage and real-time retrieval of a large amount of data and metadata, and supports flexible changes in data models by storing metadata in the JSON format, which allows a variable structure without a set schema. Moreover, the application of HDFS and HBase in KoNA and Bio-Express facilitates distributed and parallel processing of genomic big data in a high-performance computing cluster consisting of multiple computational nodes connected by the InfiniBand network. The KoNA database adopts Lucene-based Elasticsearch (version 6.3.1) [10] for data search and indexing, and Kibana (version 6.3.1) for real-time statistics visualization of a large amount of genomic metadata.

### Implementation of the KoNA web service

The KoNA web service was implemented with Spring Boot, an application framework and inversion of control container (http://www.springsource.org, version 2.2.4), and Thymeleaf, a server-side Java view template engine (version 3.0.12). All codes were developed by using Eclipse (http://www.eclipse.org), an integrated development environment for Java-based web applications. To provide stable web services, KoNA is hosted on a CentOS7 operating system with three servers: Spring Boot-based web server, an HBase server for database management, and a GBox server for fast file upload and download.

## Data availability

KoNA is available at https://www.kobic.re.kr/kona/.

## CRediT author statement

**Gunhwan Ko:** Conceptualization, Software, Writing -Original Draft. **Jae Ho Lee:** Conceptualization, Investigation, Data Curation, Writing - Original Draft, Visualization. **Young Mi Sim:** Investigation, Data Curation. **Wangho Song:** Software. **Byung-Ha Yoon:** Software, Writing - Original Draft. **Iksu Byeon:** Resources. **Bang Hyuck Lee:** Resources. **Sang-Ok Kim:** Investigation. **Jinhyuk Choi:** Software, Investigation. **Insoo Jang:** Software. **Hyerin Kim:** Methodology. **Jin Ok Yang:** Methodology. **Kiwon Jang:** Methodology. **Sora Kim:** Methodology. **Jong-Hwan Kim:** Investigation. **Jongbum Jeon:** Investigation. **Jaeeun Jung:** Methodology. **Seungwoo Hwang:** Methodology, Writing - Original Draft, Writing - Review & Editing, Supervision. **Ji-Hwan Park:** Conceptualization, Writing - Original Draft, Writing - Review & Editing, Supervision. **Pan-Gyu Kim:** Conceptualization, Supervision. **Seon-Young Kim:** Conceptualization, Writing - Original Draft, Supervision, Funding acquisition. **Byungwook Lee:** Conceptualization, Writing - Original Draft, Writing - Review & Editing, Supervision, Project administration. All authors read and approved the final manuscript.

## Acknowledgments

This research was supported by “Next-generation Genome-InfraNET” for the advancement of genome research and service” [2019M3C9A5069653] and “Construction of biological data station” [2020M3A9I6A01036057] grants from the National Research Foundation of Korea.

## Declaration of AI and AI-assisted technologies in the writing process

Nothing to disclose.

